# The resilience of the oral microbiome and lability of the hair microbiome across host environments in wild and captive lemurs

**DOI:** 10.1101/2024.09.09.612164

**Authors:** Rachel B. Burten, Richard R. Lawler, Joelisoa Ratsirarson, Jeannin Ranaivonasy, Rebecca Leduc, Ankita Bhagat, Amelia Lopez, Jason M. Kamilar

## Abstract

Microbiome diversity and composition in mammals is affected by the host’s environment and has been linked to important immune and physiological host functions, yet most of these data come from the gut microbiome. Research on the oral and hair microbiome in nonhuman primates has been far less common, and information from wild primates is even rarer. These overlooked patterns of environmental effects on microbial communities across the body may have important implications for a range of host functions. Therefore, in this study we characterized the gut, oral, and hair microbiomes across nine different captive and wild lemur species: *Eulemur collaris*, *Eulemur coronatus*, *Eulemur mongoz*, *Lemur catta*, *Microcebus griseorufus*, *Microcebus murinus*, *Propithecus coquereli*, *Propithecus verreauxi*, and *Varecia rubra*. We explored how host environment affects the microbiome diversity of these three body regions using 16S rRNA sequencing and found significant differences in microbiome composition, diversity, and environmental influence across body regions. The oral microbiome was least diverse and most resilient to different environmental effects; conversely, the hair microbiome was both most diverse and most labile. Differentially abundant bacterial taxa across oral, gut, and hair microbiota may also reflect selective regimes unique to each body region. These results emphasize the importance of accounting for body region when conducting microbiome studies.

**Importance:** An organism’s microbiome plays important roles in a wide variety of the host animal’s physiological functions, yet how these microbial communities in body regions beyond the gut are affected by the host’s environment is not clearly understood. We therefore analyzed how the oral, gut, and hair microbiomes of nine lemur species (genera *Eulemur*, *Lemur*, *Microcebus*, *Propithecus*, and *Varecia*) varied across wild and captive environments. We found that host environment affected the microbiomes of each body region in distinct ways, with the oral microbiome appearing conserved and more resilient to environmental effects, particularly compared to the diverse and widely variable hair microbiome. To our knowledge, this study is the most comprehensive multi-body-region analysis of the lemur microbiome to date. Our results demonstrate that the environment does not have a universal effect on the microbiome across body regions, but is instead mediated by body-region specific factors.

## Introduction

The importance of host microbiome diversity to the health of human and nonhuman primates has become an increasingly active area of research, but a majority of these studies focus on analyzing the gut microbiome as extrapolated from fecal samples (Gordon et al., 2005). Sampling from other regions of the body requires more invasive methods and can therefore prove challenging, especially for wild primates that do not undergo regular healthcare manipulations in captivity. Nonetheless, existing studies on microbiota from other parts of the host body demonstrate their importance within a variety of biological functions (Dewhirst et al., 2010; Greene et al., 2019; Meisel et al., 2018; Segre et al., 2010).

Information about the hair microbiome of non-human primates is particularly limited (Council et al., 2016; Segre et al., 2010; Yildirim et al., 2014). A majority of research on the primate hair microbiome focuses on humans in particular (Lousada et al., 2021; Segre et al., 2010). As primates are highly social and tactile mammals, these studies’ conclusions are likely applicable to non-human primates as well. This is because the primate hair microbiome is shaped by environmental factors, starting with the maternal microbial environment at birth when vaginal microbiota would be transferred to offspring skin and hair (Chen & Tsao, 2013). Further social transfer of hair and skin microbiota may occur in individuals that partake in allogrooming and other inter-individual tactile activities (Archie & Tung, 2015). Broader comparative research on the skin microbiome of vertebrates have identified observable phylosymbiosis of the skin microbiome at the order level (Ellison et al., 2019; Ross et al., 2018, 2019), so these taxonomic differences may not be apparent in a within-order analysis of the primate hair microbiome. This contrasts with the largest comparative study of captive non-human primate hair microbiomes to date (Kitrinos et al., 2022), which identified host species identity as the most important predictor shaping variation in the hair microbiome. Due to the constant environmental exposure of hair, its associated microbiome might be influenced by external environmental factors that differ from those impacting the gut and oral microbiota. This has been supported by multiple studies of the mammal hair/skin microbiome (Council et al. 2016; Kolodny et al. 2019; Roche et al. 2023).

Research on the oral microbiome of primates is not as sparse as hair microbiome research, though less is understood about the oral microbiome than the gut microbiome. Nonetheless, recent research on the oral microbiome in non-human primates and other vertebrates has elucidated the unique traits of this body region (Asangba et al., 2022). Asangba et al. (2022) found that the oral microbiome was compositionally similar across primate species, whereas other body regions’ microbiota clustered according to host species identity, suggesting that the oral microbiome may have a conserved function in the oral cavity. Nonetheless, these findings contrast with studies comparing the dental biofilms of anatomically modern humans, Neanderthals, and extant primates (Yates et al., 2021; Ozga et al., 2019) which identified host species-specific differences in the dental microbiome. Furthermore, a study on the preserved dental calculus microbiome of three gorilla subspecies found that host ecology shaped the oral microbiome more than geographic location or subspecies identity (Moraitou et al., 2022). These contrasting results demonstrate a need for further comparative analyses.

Host ecology and phylogeny certainly affect microbial communities across body regions (Ross et al., 2018; Council et al., 2016), but precisely *how* these factors influence microbiota depends on their location within the host’s body. Given the importance of examining microbiomes across multiple body regions within an individual, this study examines differences in the gut, oral, and hair microbiomes across nine different captive and wild lemur species: *Eulemur collaris*, *Eulemur coronatus*, *Eulemur mongoz*, *Lemur catta*, *Microcebus griseorufus*, *Microcebus murinus*, *Propithecus coquereli*, *Propithecus verreauxi*, and *Varecia rubra*. We also explore how host environment affects the microbiome diversity of different body regions. Specifically, we predict that the hair microbiome will experience larger shifts in diversity and composition across environments than the gut and oral microbiomes due to hairs’ increased contact with external substrates and microbes (Council et al., 2016). Though habitat disturbance and living in captivity will both significantly affect oral and gut microbiome alpha diversity, these body regions may cluster more by host species identity than by host environment when visualized in multivariate space due to host-specific dietary differences.

## Results

We collected oral, gut, and hair microbiome samples (Figure 1) from the nine lemur species described across five different environments: samples from *Lemur catta*, *Microcebus griseorufus*, and *Propithecus verreauxi* were collected in lower and higher disturbance areas of Beza Mahafaly Special Reserve (BMSR) in Madagascar; samples from *Eulemur collaris*, *Eulemur mongoz*, *Lemur catta*, and *Varecia rubra* were collected at Lemur Conservation Foundation (LCF) in Myakka City, Florida, USA; samples from *Lemur catta*, *Microcebus murinus*, and *Propithecus coquereli* were collected at Duke Lemur Center (DLC) in Durham, North Carolina, USA, and samples from *Eulemur coronatus* were collected from Zoo Atlanta (ZA) in Atlanta, Georgia, USA. We successfully sequenced 438 microbiome samples from five body regions across wild and captive lemurs (gut microbiome: 92, oral microbiome: 86, head hair microbiome: 88, chest hair microbiome: 89, head hair microbiome: 83. Illumina sequencing generated 26,094,555 paired-end reads after filtering for mitochondrial and chloroplast DNA and removing samples with less than 10,000 reads.

**Figure 1.**
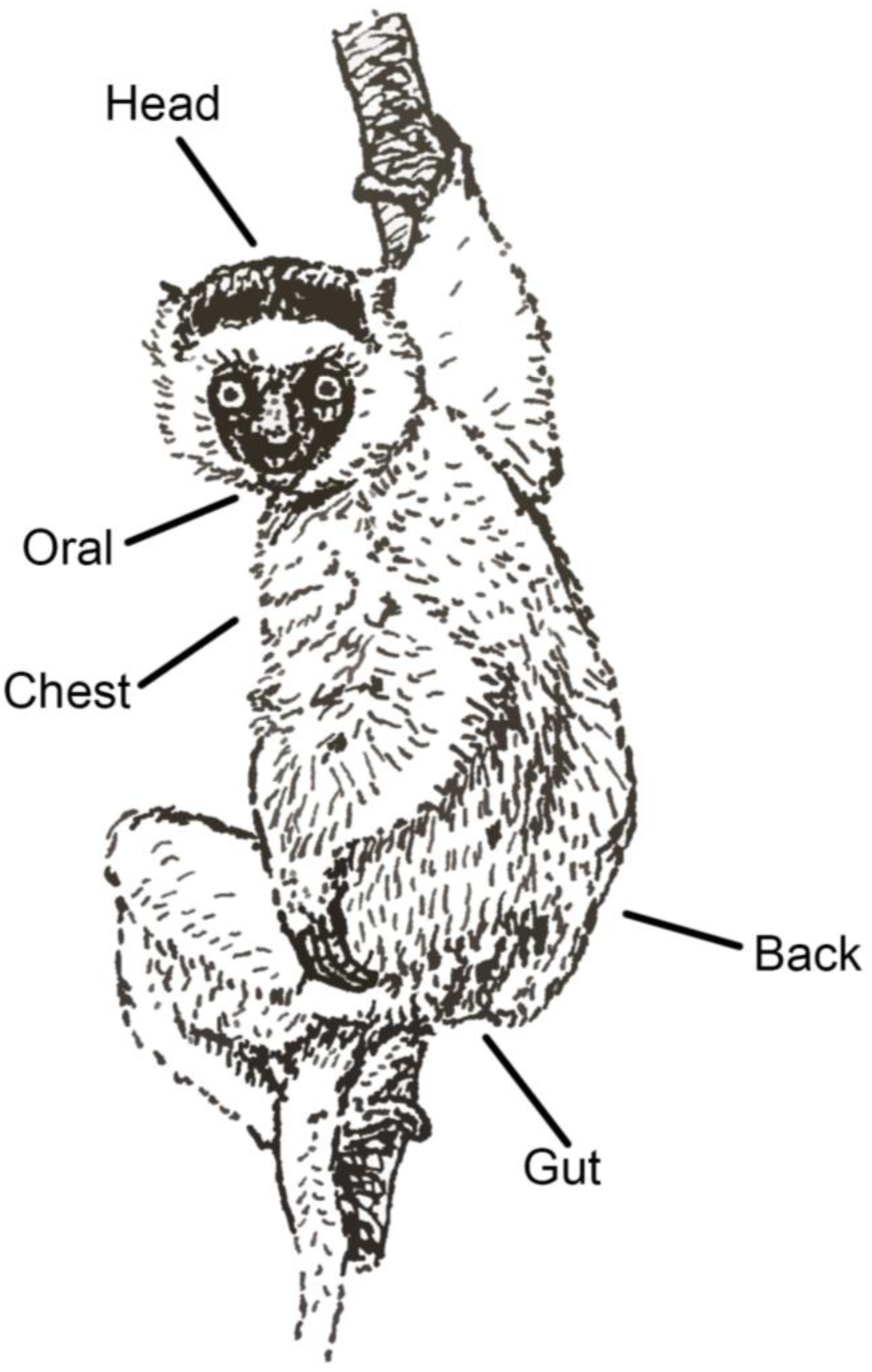
Visualization of standardized lemur sampling protocol across five body regions: head (hair), chest (hair), back (hair), oral/mouth, and gut/rectum.

### Alpha Diversity

We used three different alpha diversity indices to assess how microbiome diversity varied across body regions and host environments: Simpson, Shannon-Weaver, and Chao1. There were significant differences in alpha diversity across different body regions (Figure 2) and across different host environments for a given body region (Figure 3-5; SI Table 1). Oral microbiome sample alpha diversity varied the least across host environments, whereas gut and hair microbiome sample diversity varied more significantly. We identified significant differences between captive samples and wild samples from both high and low disturbance areas of BMSR, as well as significant differences between captive samples from different captive institutions.

**Figure 2.**
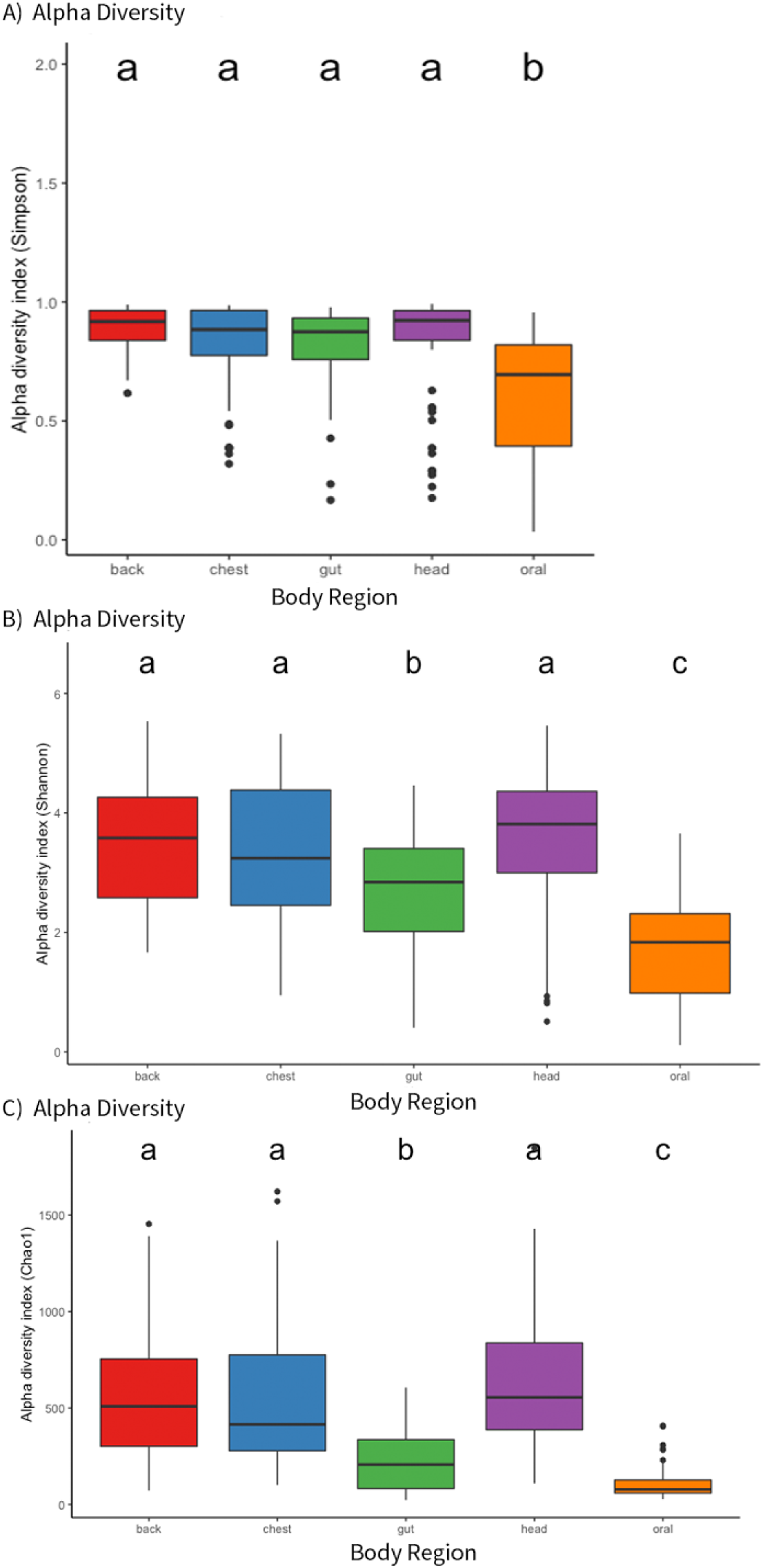
Boxplots of A) Simpson B) Shannon-Weaver and C) Chao1 alpha diversity indices for lemur microbiomes across different body regions. Letters above the boxplots symbolize statistically significant groupings across the categorical variable according to analysis of variance (ANOVA) tests followed by Tukey Honest Significant Difference (HSD) tests.

**Figure 3.**
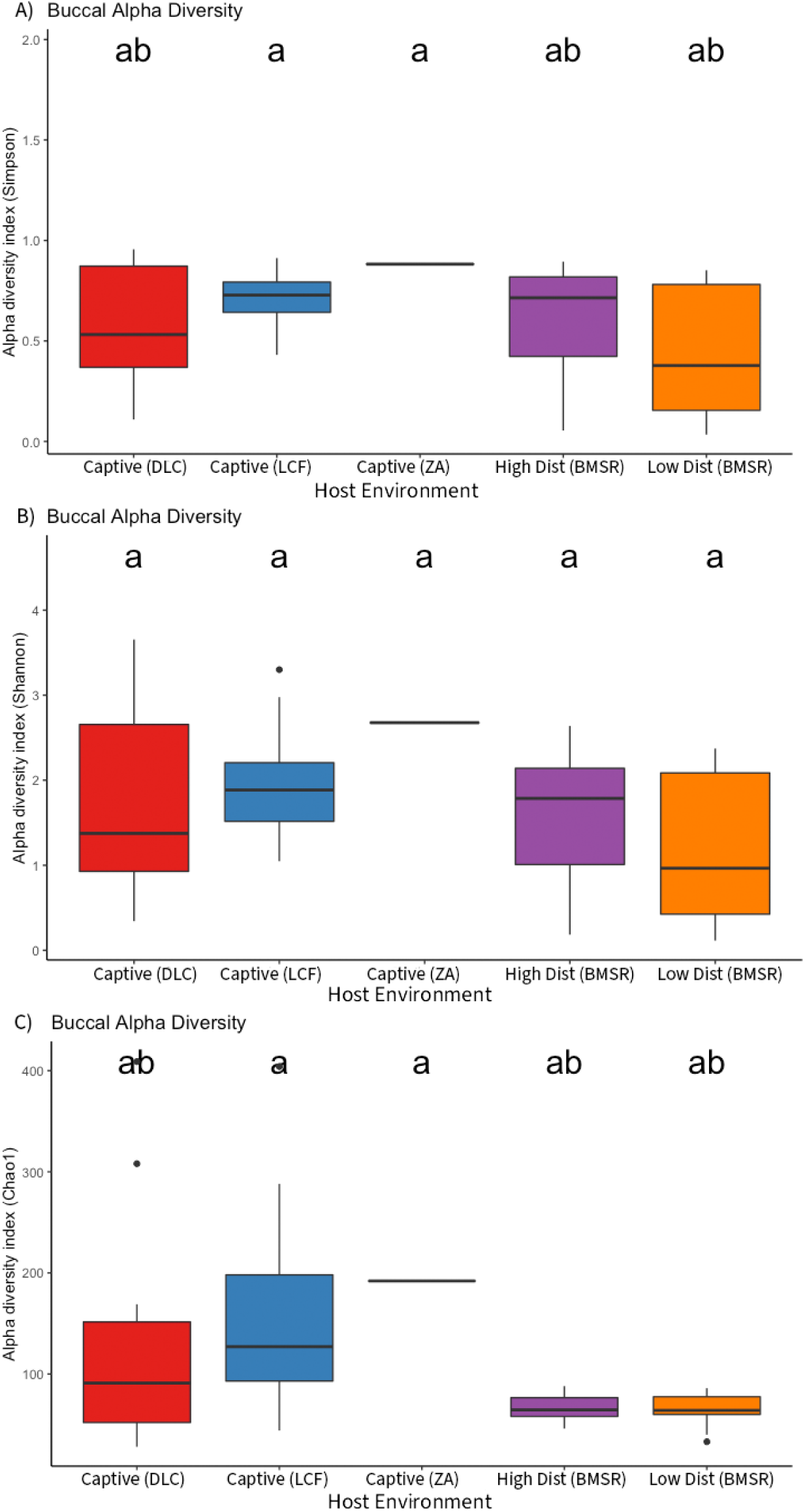
Boxplots of A) Simpson B) Shannon-Weaver and C) Chao1 alpha diversity indices for the oral microbiome of lemurs across different host environments. Letters above the boxplots symbolize statistically significant groupings across the categorical variable according to analysis of variance (ANOVA) tests followed by Tukey Honest Significant Difference (HSD) tests.

**Figure 4.**
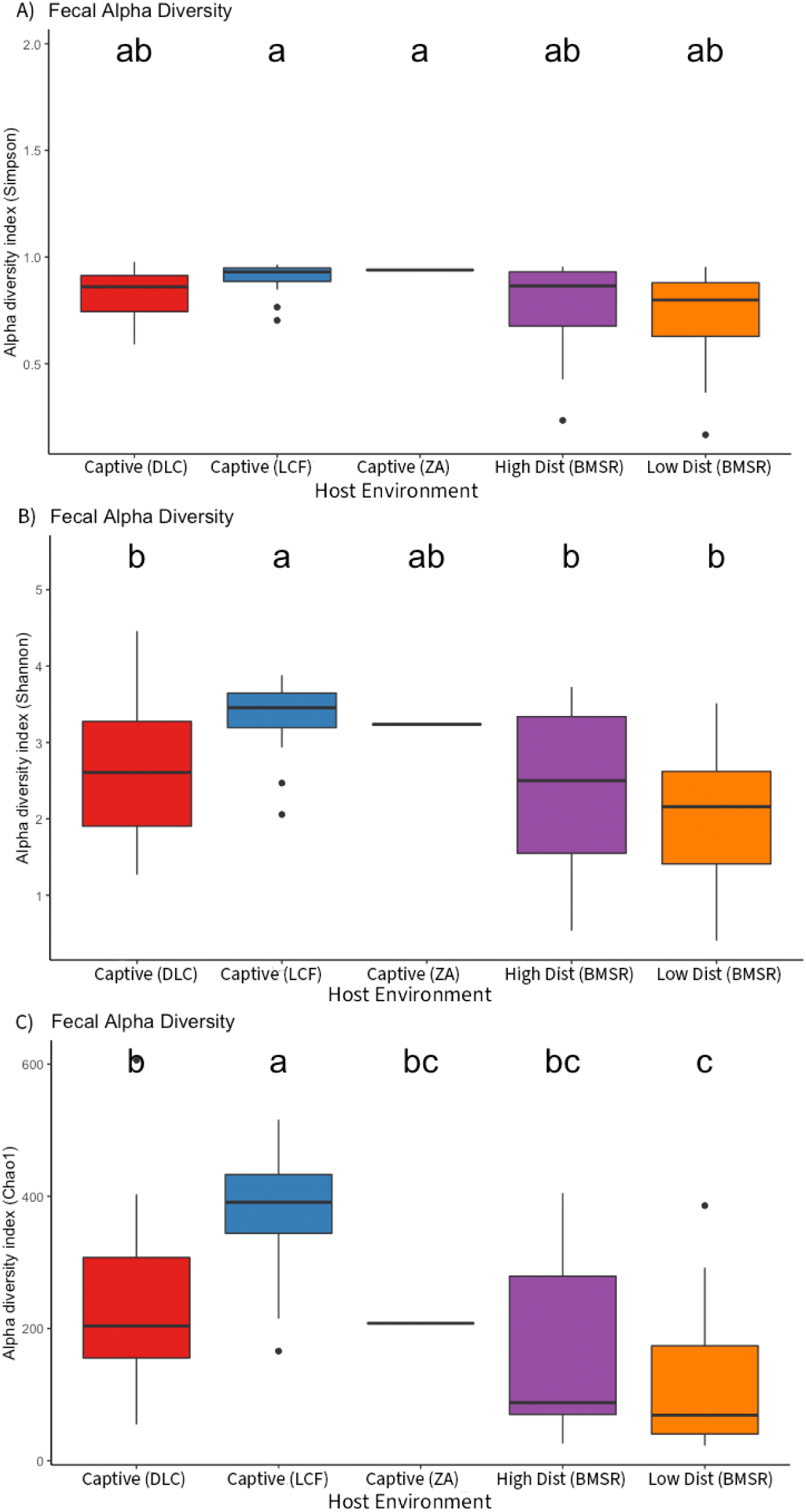
Boxplots of A) Simpson B) Shannon-Weaver and C) Chao1 alpha diversity indices for the gut microbiome of lemurs across different host environments. Letters above the boxplots symbolize statistically significant groupings across the categorical variable according to analysis of variance (ANOVA) tests followed by Tukey Honest Significant Difference (HSD) tests.

**Figure 5.**
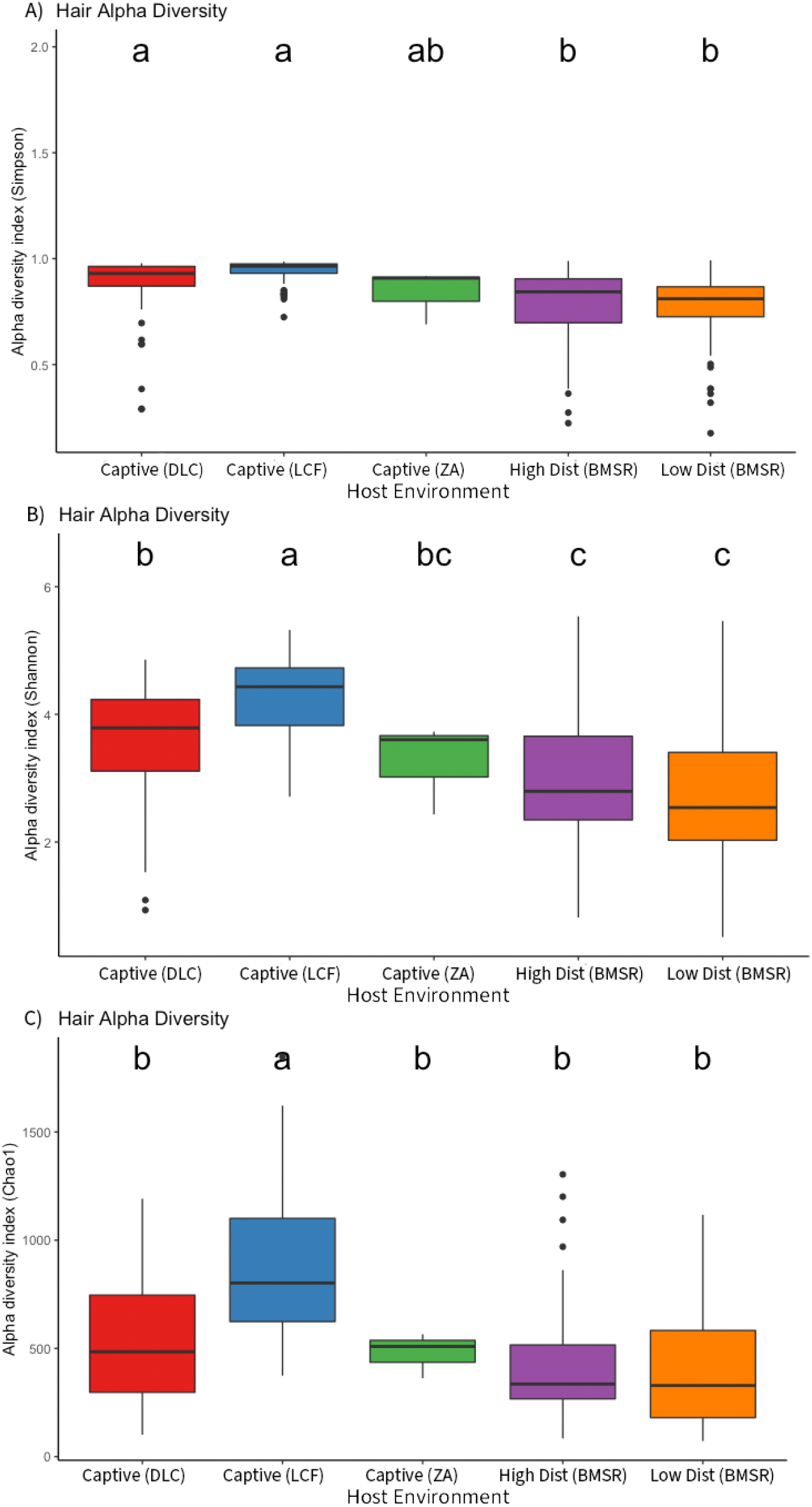
Boxplots of A) Simpson B) Shannon-Weaver and C) Chao1 alpha diversity indices for the hair microbiome of lemurs across different host environments. Letters above the boxplots symbolize statistically significant groupings across the categorical variable according to analysis of variance (ANOVA) tests followed by Tukey Honest Significant Difference (HSD) tests.

How body region affected microbial diversity also varied depending on which lemur species was being sampled (SI Table 2; SI Figures 1-4). Typically, the oral microbiome was least diverse and the hair microbiome tended to be the most diverse, with gut microbiome diversity generally in between the former and the latter. The exception to this general pattern was observed in *Microcebus* oral and gut microbiome samples, which never significantly differed from each other in diversity.

### Beta Diversity

We measured differences in microbiome composition using three beta diversity matrices: Bray-Curtis dissimilarity index, unweighted UniFrac distances, and weighted UniFrac distances. Body region significantly shaped the composition of lemur microbiomes, with oral, gut and hair microbiomes remaining compositionally distinct from each other (Figures 6, 7). Body region also explained a moderate amount of variation in microbiome composition, ranging from 6.68% to 16.51%. Furthermore, the interaction effects between body region and host species as well as body region and host environment were significant predictors of microbiome composition in the PERMANOVA tests with multiple predictor variables (SI Tables 4, 5). Differences between body regions were more apparent in Bray-Curtis multivariate space, and less apparent using unweighted UniFrac distances in multivariate space.

**Figure 6.**
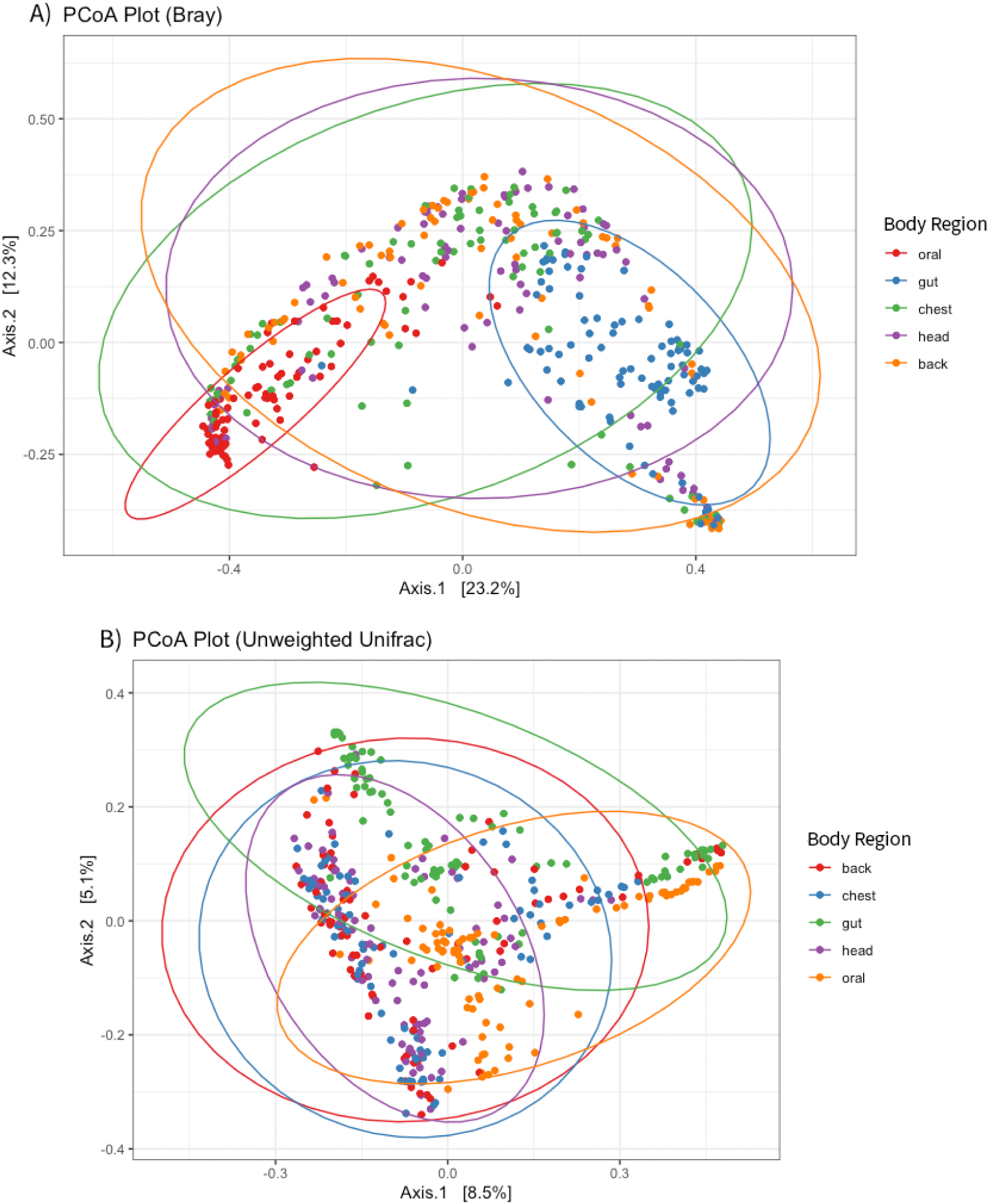
(A) Bray-Curtis Dissimilarity and (B) unweighted UniFrac Principal Coordinates Analysis (PCoA) plots of beta diversity across 438 lemur microbiome samples. The color of the points denotes the body region from which the given microbiome sample was taken, with 95% confidence intervals drawn around the five body regions: gut, oral, head hair, chest hair, and back hair.

**Figure 7.**
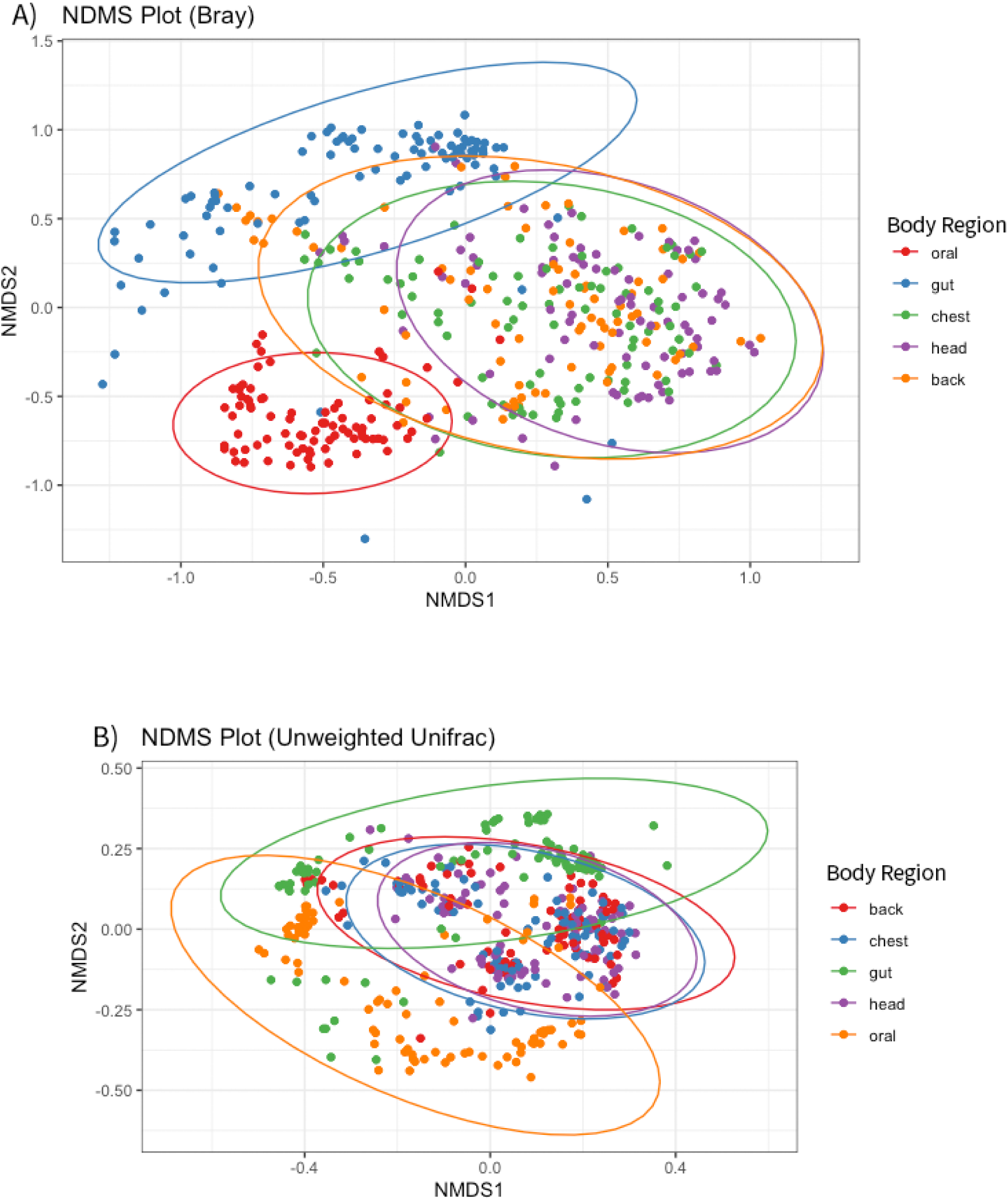
(A) Bray-Curtis Dissimilarity and (B) unweighted UniFrac non-metric multidimensional scaling (NMDS) plots of beta diversity across 438 lemur microbiome samples. The color of the points denotes the body region from which the given microbiome sample was taken, with 95% confidence intervals drawn around the five body regions: gut, oral, head hair, chest hair, and back hair.

### Linear Mixed Modeling

We ran linear mixed models (LMMs) examining how body region and other predictor variables fit the observed variation in microbiome diversity and composition, and found that body region was typically a significant and important predictor of both microbiome composition and diversity across samples. The LMMs identified significant differences in alpha diversity across hair, oral, and gut microbiomes, but less so within the three regions of hair microbiomes (Table 1). Body region had an Akaike weight of 1 for both sets of alpha diversity LMMs, and was the highest weighted predictor variable in the Shannon-Weaver diversity index LMMs.

**Table 1.** Models of best fit for each set of linear mixed models (LMMs). The eight sets of LMMs were each run with a different dependent variable measuring either alpha or beta diversity of the lemur microbiome samples. A plus (+) symbol indicates the inclusion of that fixed effect in the best-fitting model. The two best-fitting models for Bray-Curtis PCoA Axis 2 had similar Akaike weights, so both models are reported in this table.

| *Dependent Variable* | *Body Region* | *Host Environment* | *Species/Genus* | *Age* | *Sex* | *df* | *Δ AICc* | *Weight* |
| --- | --- | --- | --- | --- | --- | --- | --- | --- |
| Shannon-Weaver Alpha Diversity | + |  | + | + |  | 17 | 0 | 0.809 |
| Chao1 Alpha Diversity | + | + | + | + | + | 21 | 0 | 1 |
| Bray-Curtis Dissimilarity PCoA Axis 1 | + |  | + | + |  | 17 | 0 | 0.895 |
| Unweighted UniFrac PCoA Axis 1 | + | + | + |  |  | 17 | 0 | 0.703 |
| Weighted UniFrac PCoA Axis 1 |  | + |  |  |  | 13 | 0 | 0.988 |
| Bray-Curtis Dissimilarity PCoA Axis 2 | + | + | + |  |  | 17 | 0 | 0.512 |
| Bray-Curtis Dissimilarity PCoA Axis 2 |  | + | + |  |  | 13 | 0.1 | 0.488 |
| Unweighted UniFrac PCoA Axis 2 | + | + | + |  |  | 17 | 0 | 0.994 |
| Weighted UniFrac PCoA Axis 2 | + |  | + |  |  | 15 | 0 | 0.956 |

Body region also significantly predicted observed variation in microbiome composition for five out of the six beta diversity LMMs. For the LMMs that included Axis 1 of the Principle Coordinates Analysis (PCoA) of Bray-Curtis, Unweighted UniFrac, or Weighted UniFrac distances as the dependent variable, the gut and oral microbiomes were significantly compositionally different from the hair microbiome. The head hair microbiome and back hair microbiome also differed significantly from each other in the LMM for Axis 1 of the PCoA of Unweighted UniFrac. There was slightly more variation in the importance of body region as a predictor variable for the linear mixed models that used Axis 2 of the PCoA of Bray-Curtis, Unweighted UniFrac, and Weighted UniFrac distances, though body region was retained as a predictor variable in the best-fitting linear mixed models.

### Differential Abundance Testing

We used Analysis of Composition of Microbiomes with Bias Correction (ANCOM-BC) to examine how body region and host environment alters the abundance of bacterial and archaeal taxa. These differential abundance tests identified that there were 159 differentially abundant bacterial and archaeal taxa in the oral microbiome, 348 differentially abundant taxa in the gut microbiome, and 959 differentially abundant taxa in the hair microbiome. The ten bacterial taxa with the most significant differences (as measured by “w” score) for each body region across different host environments are detailed in Table 2. Table 3 highlights the ten bacterial taxa with the most significant differences in abundance between the head hair, chest hair, and back hair microbiome.

**Table 2.**
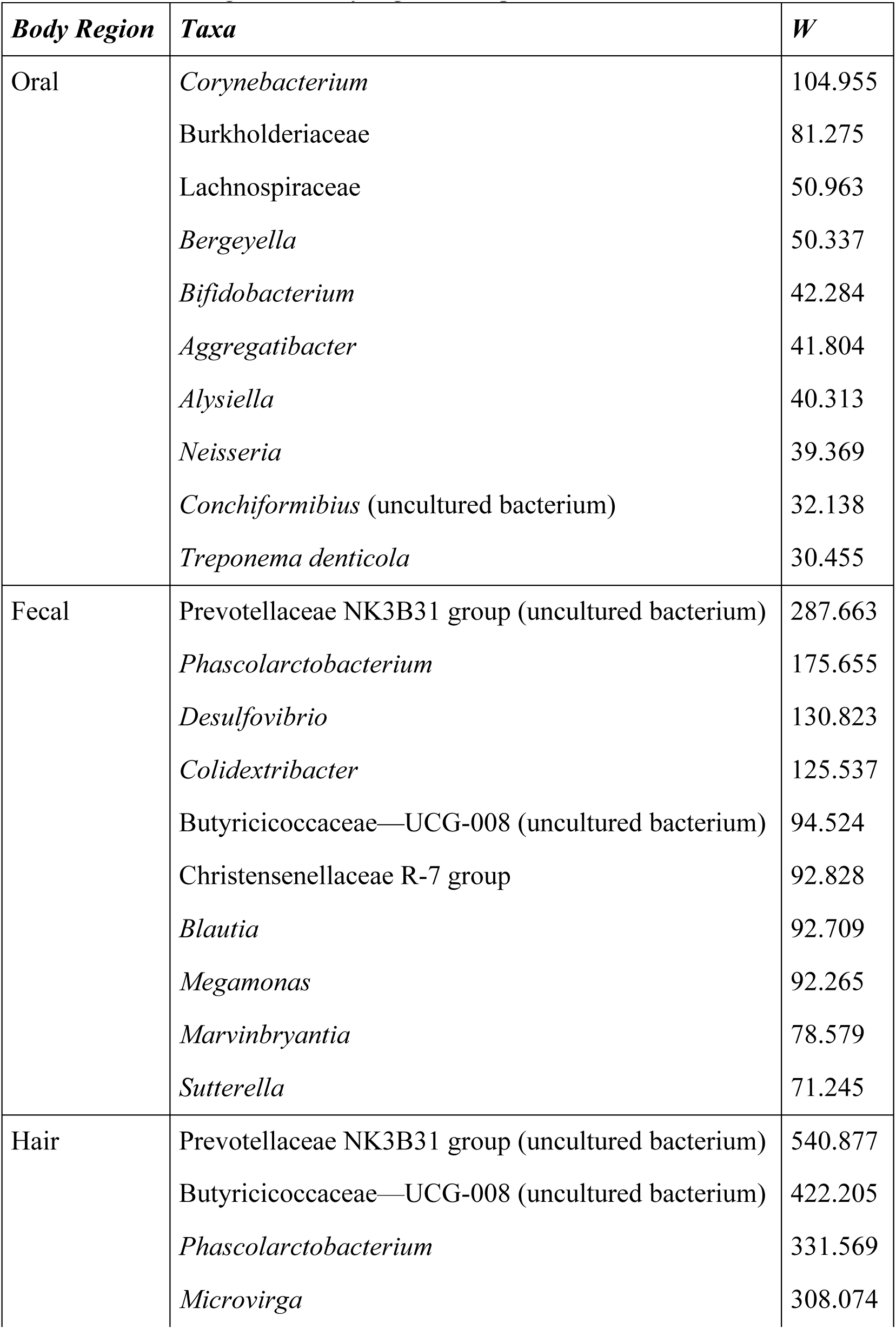
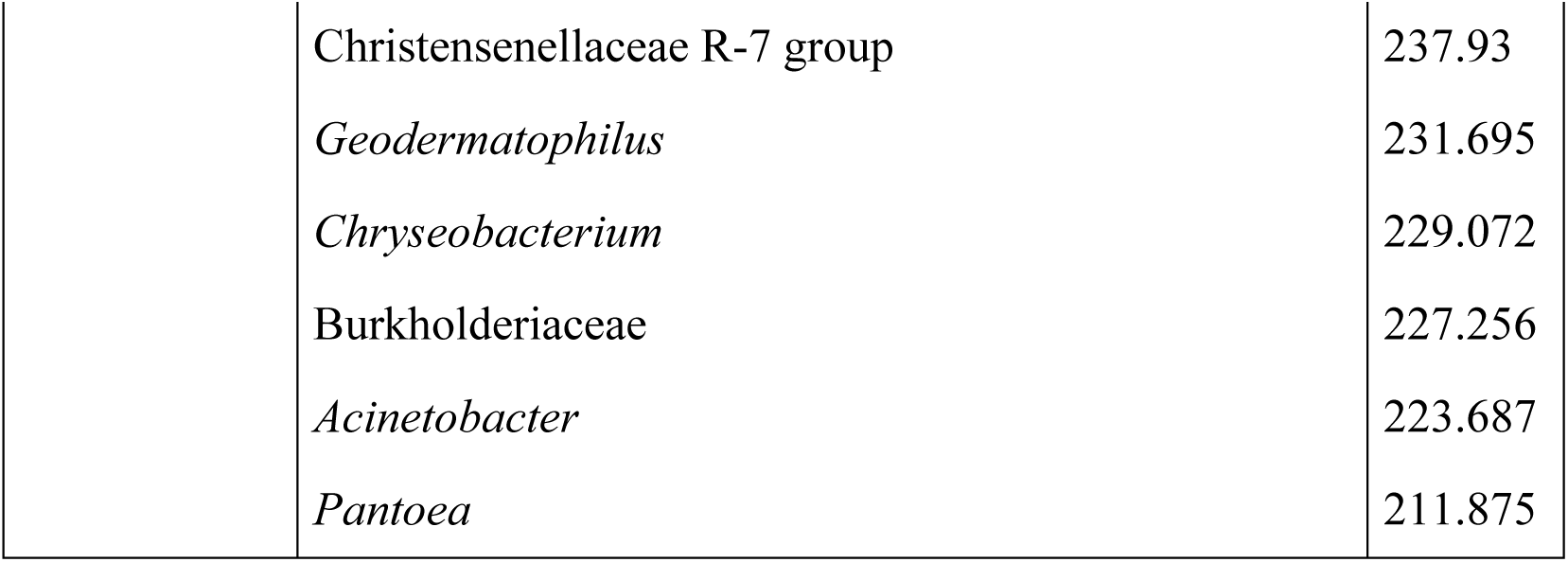
Bacterial taxa with the most significant differential abundance across host environments for each body region according to Analysis of Composition of Microbiomes with Bias Correction (ANCOM-BC). Differentially abundant taxa across host environments are reported for each of the three general body regions sampled: oral, fecal, and hair.

| *Body Region* | *Taxa* | *W* |
| --- | --- | --- |
| Oral | *Corynebacterium* | 104.955 |
|  | Burkholderiaceae | 81.275 |
|  | Lachnospiraceae | 50.963 |
|  | *Bergeyella* | 50.337 |
|  | *Bifidobacterium* | 42.284 |
|  | *Aggregatibacter* | 41.804 |
|  | *Alysiella* | 40.313 |
|  | *Neisseria* | 39.369 |
|  | *Conchiformibius* (uncultured bacterium) | 32.138 |
|  | *Treponema denticola* | 30.455 |
| Fecal | Prevotellaceae NK3B31 group (uncultured bacterium) | 287.663 |
|  | *Phascolarctobacterium* | 175.655 |
|  | *Desulfovibrio* | 130.823 |
|  | *Colidextribacter* | 125.537 |
|  | Butyricicoccaceae—UCG-008 (uncultured bacterium) | 94.524 |
|  | Christensenellaceae R-7 group | 92.828 |
|  | *Blautia* | 92.709 |
|  | *Megamonas* | 92.265 |
|  | *Marvinbryantia* | 78.579 |
|  | *Sutterella* | 71.245 |
| Hair | Prevotellaceae NK3B31 group (uncultured bacterium) | 540.877 |
|  | Butyricicoccaceae—UCG-008 (uncultured bacterium) | 422.205 |
|  | *Phascolarctobacterium* | 331.569 |
|  | *Microvirga* | 308.074 |

|  |  |
| --- | --- |
| Christensenellaceae R-7 group | 237.93 |
| *Geodermatophilus* | 231.695 |
| *Chryseobacterium* | 229.072 |
| Burkholderiaceae | 227.256 |
| *Acinetobacter* | 223.687 |
| *Pantoea* | 211.875 |

**Table 3.** Bacterial taxa with the most significant differential abundance across the head hair, chest hair, and back hair microbiomes according to Analysis of Composition of Microbiomes with Bias Correction (ANCOM-BC).

| *Body Region* | *Taxa* | *W* |
| --- | --- | --- |
| Hair | *Neisseriaceae* | 19.82 |
|  | *Altererythrobacter* | 19.663 |
|  | *Cardiobacterium* | 17.078 |
|  | *Chthoniobacter* (uncultured bacterium) | 15.805 |
|  | *Treponema maltophilum* | 15.093 |
|  | *Lautropia* sp. | 14.637 |
|  | *Bergeyella* (uncultured bacterium) | 13.598 |
|  | *Enhydrobacter* | 12.882 |
|  | WD2101 soil group (uncultured planctomycete) | 12.717 |
|  | *Flavisolibacter* | 12.607 |

## Discussion

### Differences in the Lemur Microbiome Across Body Regions

Body region is an important factor determining the composition and diversity of lemur microbiomes. Furthermore, body region-specific microhabitats and selective pressures may be generating these significant differences in the microbiome across body regions. The significant differences in the microbiome across body regions likely reflect different selective regimes acting upon these regions. The hair, oral, and gut microbiomes all vary in temperature, environmental exposure, social transmission, and the direct influence of dietary shifts. The hair microbiome had the highest diversity of any body region, which may reflect higher amounts of rare, transient bacterial taxa compared to the gut or oral microbiome. Though, there were fewer differences within different areas of the hair microbiome. Conversely, the low alpha and beta diversity of the oral microbiome may reflect a conserved microbial community that is resistant to colonization by environmental or pathogenic microbes (Asangba et al., 2022). Saliva contains proteins with antibacterial properties and physically buffers against colonization by pathogens, and oral biofilms are also incredibly resilient against colonization (Asangba et al., 2022). These differences also strongly suggest that the functional roles of microbiota vary across body regions, and existing research exploring the functional characteristics of hair, oral, and gut microbiomes supports this. The compositional differences driving the clustering of microbiome samples across body regions in multivariate space appears to be occurring mainly in closely related bacterial and archaeal taxa, which could explain why compositional differences across body regions are more apparent in the phylogeny-blind metric of beta diversity (Bray-Curtis Dissimilarity) than the phylogeny-aware metric (Unweighted UniFrac).

The hair microbial communities were consistently more diverse than either the gut or oral microbiomes. Microbial diversity and composition data are relativized: therefore, the high diversity of the hair microbiome compared to the gut and oral microbiome may reflect the presence of many rare and transient bacterial taxa. Transient environmental bacteria and archaea may be unable to colonize parts of the body that have microbiomes with conserved and specific physiological functions, which could explain the lower diversity of the oral and gut microbiomes compared to hair.

Males of multiple lemur species included in this study have scent glands on or near their chest (Greene, Bornbusch, et al., 2019), so the fact that the chest hair microbiome was generally less diverse than the back or head hair microbiome may reflect the important role of the chest hair microbiome in olfactory signaling. Thus, there may be stricter selective filtering occurring in the chest microbiome for microbes that contribute to olfactory signals. The head hair microbiome may be the most diverse across all body regions because of the frequency of grooming and social transmission of microbiota occurring on that area of the body in lemurs (Barton, 1985; Lewis, 2010). Nonetheless, there were fewer consistently significant differences in lemur hair microbiome diversity across the head, chest, and back than expected given differences in hair microstructures and exposure to grooming behaviors, environmental substrates, and other host microbial communities that would vary across areas of the hair microbiome. This aligns with previous work on the captive non-human primate hair microbiome, which found that the hair microbiome did not vary largely across different body sites (Council et al., 2016; Kitrinos et al., 2022).

Relatively little is understood about the functional properties of the hair microbiome, though research on the skin microbiome points to key roles in regulating immune responses and scent production/signaling (Council et al., 2016; Greene, Bornbusch, et al., 2019). The constant presence of low abundance transient environmental microbes could mask the signal of a resilient, conserved core hair microbiome. These results suggest that microbiome diversity across body regions is shaped by the environmental exposure and selective filtering of a given body site, with lower diversity perhaps reflecting a stronger selective regime and more conserved microbial community.

The observed significant variation in lemur microbiome composition across body regions lends further support to the important roles played by differences in environmental exposure and habitat filtering. The three hair microbiome sites were the most consistently exposed to external microbial transfer and contact with objects and other animals in the host’s environment, and in multivariate space they had the largest variation in composition. The gut microbiome and oral microbiome samples, by contrast, exhibit less intra-site variation in composition than the hair microbiome which may correspond to the increased habitat filtering and colonization resistance occurring in the oral and gut microbiomes. The fact that body region-level groupings are more apparent in multivariate space when the beta diversity metric is phylogeny-blind suggests that many of these body region differences are shaped by closely related bacterial and archaeal species across body sites. This seems to be particularly true in the hair microbiome: the spread of hair microbiome samples in multivariate space shrinks considerably when bacterial and archaeal phylogeny is taken into account. These closely related microbial species may carry out entirely different functions across body regions, but may also indicate the presence of another factor such as host species or host environment filtering for certain bacterial/archaeal taxa more strongly than body region.

Differential abundance analysis across different sites of the hair microbiome (head, chest, and back) revealed interesting differences in community membership at these sites that speak to the different environmental exposures and selective pressures present. Neisseriaceae, *Altererythrobacter*, *Chthoniobacter* (uncultured bacterium), *Lautropia* sp., WD2101 soil group (uncultured planctomycete), and *Flavisolibacter* were all higher abundance in head hair than other hair microbiomes. Neisseriaceae and *Lautropia* sp. are closely associated with the oral cavity (Lim et al., 2019; Liu et al., 2015). Conversely, *Altererythrobacter*, *Chthoniobacter* (uncultured bacterium), WD2101 soil group (uncultured planctomycete), and *Flavisolibacter* are commonly found in environmental and soil microbial communities (Dedysh et al., 2021; Lee et al., 2016; Sangwan et al., 2004; Xu et al., 2020). *Enhydrobacter* has been associated with moisture content and sebum production on human skin (Kim et al., 2021; Mukherjee et al., 2016), and was most abundant in the chest microbiome and lowest in the head hair microbiome. The generally high abundance of oral and soil-associated microbiota on the head hair microbiome may reflect the high rates of grooming with a tooth comb that occurs around the head in lemurs (Barton, 1985; Lewis, 2010), as well as a high level of exposure to environmental substrates such as soil and plants. The enriched abundance of *Enhydrobacter* in the lemur chest microbiome may reflect increased oil production on the lemur chest for olfactory signaling, and warrants further exploration.

### The Relationship Between the Microbiome and Host Environment is Mediated by Body Region

As predicted, body region mediated the effect of the environment on the lemur microbiome. The interaction effect between host environment and body region was also a significant determinant of microbiome composition, though it typically only explained a moderate amount of the observed compositional variance. The oral microbiome in particular seemed least affected by differences in host environment, whereas the hair microbiome appeared to be the most significantly shaped by these differences. The gut microbiome was generally intermediate between the two. Additionally, both body region and host environment were included in the linear mixed model of best fit for half of all LMMS runs. We identified statistically significant differences in microbiome alpha and beta diversity across body regions using LMMs, though these same LMMs only identified statistically significant differences in microbiome beta diversity—but not alpha diversity—across different host environments. Therefore, body region may have a larger effect on the within-sample diversity of the lemur microbiome than host environment. These results emphasize the importance of considering body region sampled when conducting microbiome studies, as the influence of a given external factor on the microbiome may vary significantly by body region.

Across all body regions, average microbiome alpha diversity was highest in captive samples and lowest in wild samples, though these differences were smaller and mostly insignificant in oral samples and more prominent in hair samples. These results are not in agreement with many previous studies on the effects of captivity on the primate microbiome, but do align with more recent research on captive and wild lemur populations (Bornbusch et al., 2022; Greene et al., 2021). Furthermore, these results align with our predictions that the hair microbiome would be most vulnerable to external environmental input. The fact that the oral microbiome was more resistant to environmental effects than the gut microbiome was not predicted, but is supported by some existing research on the particularly conserved nature of the oral microbiome (Asangba et al., 2022).

The results of tests examining variation in microbiome diversity and composition across different body regions were surprisingly congruent with each other. Analysis of variance tests exploring how alpha diversity varies across host environments for a given body region revealed few significant differences in oral microbiome diversity across environments. Similarly, the composition of all oral microbiome samples varied less from each other in multivariate space than any other body region. The relatively conserved nature of the oral microbiome across different environments suggests that the host’s external environment is not the most important factor structuring the oral microbiome, and that the oral microbiome is more resistant to environmental perturbation than other body regions. This may be due to key functions carried out by the oral microbiome that remain unchanged even in light of dietary differences across environments (Asangba et al,. 2022). The oral microbiome results align with Asangba et al.’s (2022) findings on the conserved nature of the oral microbiome, but differ from other recent studies which found that host species identity and/or environment were important determinants of the oral microbial community (Yates et al., 2021; Moraitou et al., 2022; Ozga et al., 2019). However, these contrasting studies specifically analyzed the microbiome of preserved dental calculus as opposed to the general oral cavity sampling conducted by Asangba et al. (2022) and the buccal/general oral sampling carried out in this study. The difference in results from dental calculus microbiome studies and general/buccal oral microbiome studies highlights how microhabitat variation within the oral cavity could mean multiple different microbial communities with unique selective regimes exist there. Biofilms are a functionally important, common microbial structure within the oral cavity that may also be more resistant to colonization of environmentally-introduced bacteria (Mosaddad et al., 2019; Motta et al., 2021).

The gut microbiome varied more across host environments than the oral microbiome, but was less labile than the hair microbiome. This indicates that host environment has a moderate and detectable impact on gut microbiome diversity, likely through dietary changes and passive ingestion of environment-associated microbiota. From a functional perspective, it is adaptive for the gut microbiome to be more labile and differentiate across host environments. The gut microbiome in lemurs is essential for breaking down, fermenting and accessing macro/micronutrients from ingested food (Amato et al., 2016), so the gut as a microhabitat would select for microbes that can best process the specific items being ingested. These food items, of course, would vary across host environments. At the same time, the vital physiological functions of the gut microbiome and/or functional flexibility of certain microbes may result in the conservation of some microbial taxa across environments, regardless of host diet. This balance can be observed when visualizing gut microbiome sample composition in multivariate space: while there is more variation than oral samples, there is less variation across both axes compared to hair microbiome samples.

The significant differences in hair microbiome diversity across host environments paired with the wide variety in hair microbiome composition suggests that the hair microbiome is largely influenced by external factors. This is similar to multiple studies on the hair microbiome of bat colonies (Kolodny et al., 2019; Lemieux-Labonté et al., 2016)) which identified shared environments and social group associations as more important for structuring the hair microbiome than individual ID or host species identity. Given the strong effect of host environment on hair microbiome diversity and composition, we suggest that the hair microbiome can provide key insights into how the host interacts with their environment and their social networks (Archie & Tung, 2015; Kolodny et al., 2019).

Differential abundance testing elucidated how these body regions may be specifically impacted by host environment. In the oral microbiome, *Corynebacterium*, Burkholderiaceae, *Bergeyella*, *Aggregatibacter,* and *Treponema denticola* were significantly enriched in captive lemurs compared to wild lemurs. *Corynebacterium* and *Treponema* species are both expected members of the mammal microbiome, but have been respectively associated with human diseases and pectin degradation in fruits (Greene, Bornbusch, et al., 2019). The higher abundance of these taxa in captivity thus lend support to the homogenizing and Westernizing nature of captivity on the microbiome. Lachnospiraceae, Bifidobacterium, *Neisseria*, and *Conchiformibius* were present in significantly higher abundances in low disturbance habitat from BMSR compared to high disturbance areas or captivity. Many of these bacterial taxa are associated with important functions in the primate microbiome, such as production of the short-chain fatty acid butyrate, reduction of inflammation, and immune responses (Aho et al., 2020; Henrick et al., 2018; Jeong et al., 2021; Vital et al., 2017). The enrichment of these microbes in wild lemur oral microbiome could suggest a challenging host environment in which individuals need to rely more heavily on SCFA production within their microbiomes to derive energy from their diet, and whose immune systems are more frequently challenged.

Soil-associated bacteria were enriched in wild lemur hair microbiomes compared to captive hair microbiomes, suggesting that the hair microbiome can serve as an excellent indicator of the environmental microbes to which the host lemur is exposed. Rhizobiaceae was significantly enriched in the *M. griseorufus*, *P. verreauxi*, and (to a lesser degree) *V. rubra* hair microbiome. Rhizobiaceae is a plant and soil-associated bacterial taxa, but has been found in studies of the skin microbiome in both humans and Malagasy frogs (Bletz et al., 2017; Manus et al., 2023). Its presence in the *P. verreauxi* and *M. griseorufus* microbiomes (the two lemur species with hair microbiome samples collected in the wild) could be a better indicator of environmental impact on the hair microbiome rather than host species-specific differences.

Similarly, *Microvirga* and *Geodermatophilus* were significantly less abundant in the captive lemur microbiome, and enriched in the low-disturbance areas of BMSR. Both of these genera are soil-associated, reinforcing the importance of host species-specific environmental interaction (such as terrestrial behavior) in shaping the hair/skin microbiome (Montero-Calasanz, 2023; Msaddak et al., 2017).

### Future Directions

Differential abundance tests within the hair microbiome suggest that differences detected across hair microbiomes are likely determined by each hair site’s specific relationship to allogrooming/social behaviors and olfactory signaling. However, the density, length, color, and texture of hair varies across these body regions in lemurs and may be shaping the hair microbiome in ways we did not explicitly test for in this study. Future analyses that pairs investigation into hair microstructure variation with microbiome diversity and composition could elucidate the role of hair morphology in the hair microbiome, as has been done with paired analysis of lemur gut morphology and the gut microbiome (Greene et al., 2023).

The results of this study demonstrate that the environment does not have a universal, identical effect on the microbiome across body regions, but rather is mediated by body-region specific factors. Future studies exploring how the environment is altering the microbiome of a non-human primate should strongly consider incorporated multiple body sites to get a more complete answer to this question. If only one body region is examined, then the unique influence of this body region on the microbiome-environment relationship should be taken into account.

For example, the hair microbiome may be a particularly excellent indicator of the environmental microbes with which the host comes into contact, whereas changes to the oral and gut microbiome may be better indicators of the functional differences across environments.

## Methods

### Study Sites and Body Regions

We collected microbiome samples using a standardized protocol across five body regions in lemur species at three captive institutions in the United States and one wild setting in southwest Madagascar. We chose to collect samples from the oral cavity, rectum, head hair, chest hair, and back hair microbiome because these five body sites hypothetically have extremely different microenvironments and require different physiological functions from their microbial communities (Asangba et al., 2022). Hair length, texture, and density also vary between these regions on the lemur body, which could further drive variation in the microbiome. All wild lemur samples were obtained from individuals at Beza Mahafaly Special Reserve (BMSR) in southwest Madagascar in August-September 2018 and August-September 2019. Captive lemur sampling was conducted at Duke Lemur Center (DLC), Lemur Conservation Foundation (LCF), and Zoo Atlanta (ZA). Samples from DLC were collected between April 2022 and July 2022; samples from LCF were collected between May 2022 and December 2022; and samples from ZA were collected between May 2022 and October 2022. All captures and sampling procedures carried out in this study received IACUC approval prior to sample collection (See Supplemental Materials for detailed sampling and capture methods).

### Sample Preparation, Processing, and Sequencing

Our genomic protocols followed a modified version of the Earth Microbiome Project (Caparoso et al., 2012). We first extracted microbial DNA from all samples using the PureLink Microbiome DNA Purification Kit (Thermo Fisher Scientific) and following the kit’s oral swab protocol with a modification of incubation for 10 minutes at 95° Celsius instead of 65° Celsius. Secondly, we completed PCR amplifications targeting the 515-806 V4 hypervariable region of the 16S ribosomal RNA gene (approximately 390 bp) using primers with Illumina adapters and unique GoLay barcodes to allow multiplexing of pooled samples. DNA library size and quality was further examined at the Genomics Resource Laboratory at the University of Massachusetts Amherst via a high sensitivity DNA assay on the Agilent 2100 Bioanalyzer system. Finally, sample sequencing was completed across three sequencing runs at the Genomics Resource Laboratory on MiSeq Illumina platforms with V3 chemistry, generating 201bp reads.

### Data Analysis

We used the Massachusetts Green High-Performance Computing Cluster (MGHPCC) to process the amplicon sequences within the Quantitative Insights into Microbial Ecology (QIIME 2) pipeline (Bolyen et al., 2019). We ran the DADA2 (Callahan et al., 2017) plugin within QIIME 2 to identify amplicon sequence variants (ASVs) and to correct and remove sequencing errors and chimeric sequences. Subsequently, we used the naive Bayesian classifier method trained on SILVA 138 515F/806R reference sequences (Quast et al., 2012) clustered at 99% similarity to assign taxonomy to the ASVs. Samples with read depths below 10,000 were removed from further analyses. We used scaling with ranked subsampling (SRS) to normalize sample reads for downstream analyses via the SRS package in R (Beule and Karlovsky, 2020). Samples were scaled to the read depth of the sample with the fewest reads: 11,870.

We calculated microbial alpha diversity for each sample using the Shannon-Weaver and Simpson observed richness diversity indices (Simpson, 1949; Lemos et al., 2011), as well as the Chao1 estimated richness diversity index (Chao, 1984). We used the SRS-normalized ASV counts for the Shannon-Weaver and Simpson indices, and the raw ASV counts for the Chao1 index. We calculated beta diversity using the Bray-Curtis dissimilarity index (Bray & Curtis, 1957) as well as weighted and unweighted UniFrac distances (Lozupone & Knight, 2005).

Subsequently, we used the Bray-Curtis and unweighted UniFrac distances in principal coordinates analysis (PCoA) plots and non-metric multidimensional scaling (NMDS) plots that included all samples to visualize beta diversity (McMurdie & Holmes, 2013).

We ran three sets of analysis of variance (ANOVA) tests and post-hoc Tukey Honest Significant Difference (HSD) tests to examine 1) if microbiome alpha diversity differs across body regions, 2) if alpha diversity differs across host environments for a given body region, and 3) if alpha diversity differs across body regions within a given host species/genus identity. We used the *vegan* and *agricolae* packages in R (de Mendiburu, 2019; Oksanen et al., 2013) to run the ANOVA tests and the post-hoc Tukey HSD tests, and conducted these tests with the 1) Shannon-Weaver, 2) Simpson, and 3) Chao1 diversity indices alternatively serving as the dependent variable.

Subsequently, we used permutational multivariate analysis of variance tests (PERMANOVA) to explore how the composition of microbiome samples differs across different body regions (Oksanen et al., 2013). We ran three sets of PERMANOVA tests with different predictors: 1) body region only as the categorical predictor variable, 2) species, host environment, body region, and the interactions between these groups as the group predictor variables; and 3) species, host environment, body region, the interaction between body region and host environment, and the interaction between body region and species as the group predictor variables. To examine how different dissimilarity measurements may affect these results, We used Bray-Curtis, unweighted UniFrac, and weighted UniFrac distance matrices as the measures of community composition in these PERMANOVA tests. The number of permutations was set to 1000 for each PERMANOVA test.

Next, we ran linear mixed models using the *lme4* (Bates et al., 2014) and *lmerTest* (Kuznetsova et al., 2017) packages in R to analyze how body region affects microbiome diversity and composition and to account for potential covariation among predictor variables. We also used these LMMs to understand which factors best explain variation in microbiome diversity and composition, and to this end included host species identity, host environment, host age, and host sex as fixed effects in these models, with individual ID set as a random effect.

These LMMs were run with eight different dependent variables: Axis 1 and Axis 2 of principal coordinates analysis generated from Bray-Curtis Dissimilarity, unweighted UniFrac distances, and weighted UniFrac distances were all used to examine how microbiome composition differs across body regions and other factors of interest. Additionally, Chao1 and Shannon-Weaver indices were used to examine how microbiome diversity is affected by body region and the other independent variables. We subsequently used the MuMIn package in R (Barton, 2012) for model selection to determine which subsets of independent variables best account for variation in microbiome composition and diversity. We evaluated the models of best fit using the corrected Akaike’s Information Criterion (AICc) and Akaike weight.

We were also interested in determining how body region and host environment interact to shape the differential abundance of bacterial and archaeal taxa in the lemur microbiome. We therefore used Analysis of Compositions of Microbiomes with Bias Correction (ANCOM-BC) in the *ANCOMBC* package (Lin and Peddada, 2020) to explore how host environment affects the abundance of bacteria and archaea across different body regions (oral, fecal, and hair). ANCOM-BC allows for differential abundance testing across interaction effects between independent variables, so hair microbiome samples were further examined to understand how both specific regions of the hair microbiome and host environment shape the differential abundance of bacteria and archaea.

## Acknowledgements

This research was funded by the University of Massachusetts Amherst Predissertation Grant (RBB), Dissertation Fieldwork Grant (RBB), and Natural History Collections Summer Scholarship (RBB), as well the National Science Foundation Graduate Research Fellowship Program (RBB), the National Science Foundation Doctoral Dissertation Improvement Grant (BCS #2120509) (RBB and JMK), and the Leakey Foundation (RBB).

We thank Courtney Babbitt, Brenda Bradley, Kristen DeAngelis, and Toni Lyn Morelli for their invaluable comments and suggestions which greatly improved the manuscript.

